# Intravital imaging reveals cell cycle-dependent satellite cell migration during muscle regeneration

**DOI:** 10.1101/2020.05.22.111138

**Authors:** Yumi Konagaya, Kanako Takakura, Maina Sogabe, Anjali Bisaria, Chad Liu, Tobias Meyer, Atsuko Sehara-Fujisawa, Michiyuki Matsuda, Kenta Terai

## Abstract

During muscle regeneration, extracellular signal-regulated kinase (ERK) promotes both proliferation and migration. However, the relationship between proliferation and migration is poorly understood in this context. To elucidate this complex relationship on a physiological level, we established an intravital imaging system for measuring ERK activity, migration speed, and cell-cycle phases in mouse muscle satellite cells. We found that *in vivo*, ERK was maximally activated in satellite cells two days after injury, and this is then followed by increases in cell number and motility. With limited effects of immediate ERK activity on migration, we hypothesized that ERK increases migration speed in the later phase by promoting cell-cycle progression. Our cell-cycle analysis further revealed that in satellite cells, ERK activity is critical for the G1/S transition, and cells migrate more rapidly in the S/G2 phase three days after injury. Finally, migration speed of satellite cells was suppressed after CDK1/2, but not CDK1, inhibitor treatment, demonstrating a critical role of CDK2 in satellite cell migration. Overall, our study demonstrates that in satellite cells, the ERK-CDK2 axis not only promotes the G1/S transition, but also migration speed, which may provide a novel mechanism for efficient muscle regeneration.

## Introduction

To efficiently regenerate skeletal muscles, the right cells to be at the right place at the right time. This coordinated process is dependent on muscle stem cells, or so called satellite cells, that reside quiescent in uninjured muscles (Yin et al., 2013); (Ceafalan et al., 2014); (Tedesco et al., 2010). Upon injury, activated satellite cells start proliferation and differentiate into myoblasts. Myoblasts proliferate, migrate to the site of injury, and then differentiate into myofibers, completing the regeneration process. A subpopulation of satellite cells undergoes self-renewal to restore the pool of quiescent satellite cells. Recent studies have indicated that dysfunction of satellite cells can contribute to age-associated muscle diseases and influence genetic disorders such as Duchenne muscular dystrophy (DMD) (Blau et al., 2015); (Sousa-Victor et al., 2015); (Almada and Wagers, 2016).

Several myogenic transcription factors are sequentially activated to restore muscle structure and function after injury. Satellite cells express the transcription factor paired box 7 (PAX7), which is essential for satellite cell survival and muscle regeneration (Seale et al., 2000); (Oustanina et al., 2004); (Kuang et al., 2006). Satellite cell activation is characterized by the expression of myogenic determination protein (MYOD) and myogenic factor 5 (MYF5). The differentiation of myoblasts involves the downregulation of PAX7 and the expression of myogenin (MYOG) (Ceafalan et al., 2014); (Tedesco et al., 2010).

The relationship between proliferation and migration is complex and context-dependent, and has been mostly studied in tumorigenesis and development. Historically, cancer cell proliferation and migration were considered to be mutually exclusive in time and space, which is often referred to as the “go or grow” hypothesis (Giese et al., 1996); (Corcoran et al., 2003); (Garay et al., 2013). This hypothesis is corroborated by reports showing that tumor cells in the G0/G1 phase migrate more vigorously than in the S/G2/M phase (Bouchard et al., 2013); (Yano et al., 2014). However, several lines of evidence indicate that tumor cells can migrate faster in the S/G2/M phase compared to G0/G1 phase (Kagawa et al., 2013); (Haass et al., 2014). In development, neural crest cells in fish and avian embryo migrate faster in S phase (Burstyn-Cohen and Kalcheim, 2002); (Rajan et al., 2018). And during mouse cerebral cortex development, nuclei of neural progenitors in the ventricular zone migrate more vigorously in the S/G2/M phase than in G1 (Sakaue-Sawano et al., 2008). It is thus likely that the relationship between proliferation and migration depends on the cells, tissues, and the surrounding environment, and much remains unknown in other physiological context such as muscle regeneration.

Extracellular signal-regulated kinase (ERK) signaling pathway has been suggested to play crucial roles in muscle regeneration. Previous studies have shown that ERK1/2 promotes myoblast proliferation and migration *in vitro* (Suzuki et al., 2000); (Jones et al., 2001). In addition, ERK1/2 has also been reported to be important for muscle differentiation *in vitro* (Rommel et al., 1999); (Yokoyama et al., 2007); (Koyama et al., 2008) and *in vivo* (Michailovici et al., 2014). *Erk1-/-* mutant mice have 40% less quiescent satellite cells compared to control (Le Grand et al., 2012), further emphasizing the importance of ERK signaling in satellite cells. In many of these reports, fibroblast growth factor (FGF) acts upstream of the ERK signaling pathway. The significance of FGF is highlighted by a muscle regeneration defect in *FGF6-/-* mutant mice (Floss et al., 1997), severe muscular dystrophy in *FGF2-/-/FGF6-/-/mdx* mutant mice (Neuhaus et al., 2003), and enhanced wound repair by the delivery of FGF2 (Doukas et al., 2002). However, when and to what extent ERK plays its critical roles for muscle regeneration remains poorly understood.

Intravital imaging by two-photon microscopy is becoming a powerful technique to study the complexity of biological events in living tissues including skeletal muscle (Pittet and Weissleder, 2011); (Nobis et al., 2018). For example, Webster et al. developed an intravital imaging technique to observe cells labeled with Pax7-CreERT2 in living mice (Webster et al., 2016). They demonstrated that extracellular matrix (ECM) remnants guide the direction of migration and division plane. Another intravital imaging technique developed by Mercier et al. revealed that single fibers contraction occurs spontaneously and independently of neighboring fibers within the same muscle (Lau et al., 2016). More recently, Hotta et al. revealed that the temporal profile of microvascular hyperpermeability to be related to that of eccentric contraction-induced skeletal muscle injury (Hotta et al., 2018). Thus, intravital imaging provides the information on biological events including cell division, cell migration, myofiber contraction, and vascular permeability, which could never be obtained without intravital imaging.

To further understand the role of ERK signaling and how cell migration is affected by cell-cycle modulations during muscle regeneration *in vivo*, we established an intravital imaging technique to observe live mouse muscle regeneration. We incorporated in this imaging system a Förster/fluorescence resonance energy transfer (FRET) biosensor that measures ERK and activity and a fluorescent reporter that indicates cell cycle. With this intravital imaging platform, we found that ERK promotes the G1/S phase transition and that satellite cells migrate faster in the S/G2 phase. Moreover, our data suggests that CDK2 is responsible for promoting migration speed of satellite cells. In summary, our study clarifies the cell cycle-dependent migration of satellite cells *in vivo*, and may provide a novel mechanism of efficient tissue regeneration.

## Results

### ERK is activated during muscle regeneration

Satellite cell proliferation and migration have been reported to be essential for muscle regeneration. To investigate the relationship between satellite cell proliferation and migration, we focused on ERK, which has been reported to promote both myoblast proliferation (Jones et al., 2001) and migration (Suzuki et al., 2000) *in vitro*. To study ERK activity in living tissues, we used a previously developed R26R-EKAREV mice strain that ubiquitously expressed a floxed FRET biosensor for monitoring ERK activity, EKAREV (Konishi et al., 2018). We crossed the R26R-EKAREV mice Pax7-CreERT2 mice (Lepper et al., 2009) to generate R26R-EKAREV/Pax7-CreERT2 mice (Fig. 1A). Skeletal muscle damage was induced by cardiotoxin injection, and then live imaged under an upright microscope via imaging window (Fig. 1B and 1C) (Takaoka et al., 2016). After Cre-mediated recombination induced by tamoxifen, R26R-EKAREV/Pax7-CreERT2 mice express a FRET biosensor for ERK, in the nucleus of Pax7 lineage cells, hereinafter referred to as satellite cells (green cells and pseudo-colored cells in Fig. 1D). Cells that were not recombined, i.e., myofibers, expressed a large Stokes shift fluorescent protein, tdKeima. We confirmed that tdKeima was expressed ubiquitously in muscle fibers before injury (magenta cells in Fig. 1D). Marked reduction in the number of tdKeima-expressing cells was observed between 0 and 2 days post injury (dpi) (Fig. 1D and 1E). The nuclear density of satellite cells was measured from the z-stack images of skeletal muscle, and assessed by a multiple contrast method, Scheffe’s F-test. The nuclear density was increased by 3.2 fold from 2 to 3 dpi, indicating the proliferation of satellite cells.

**Figure 1.**
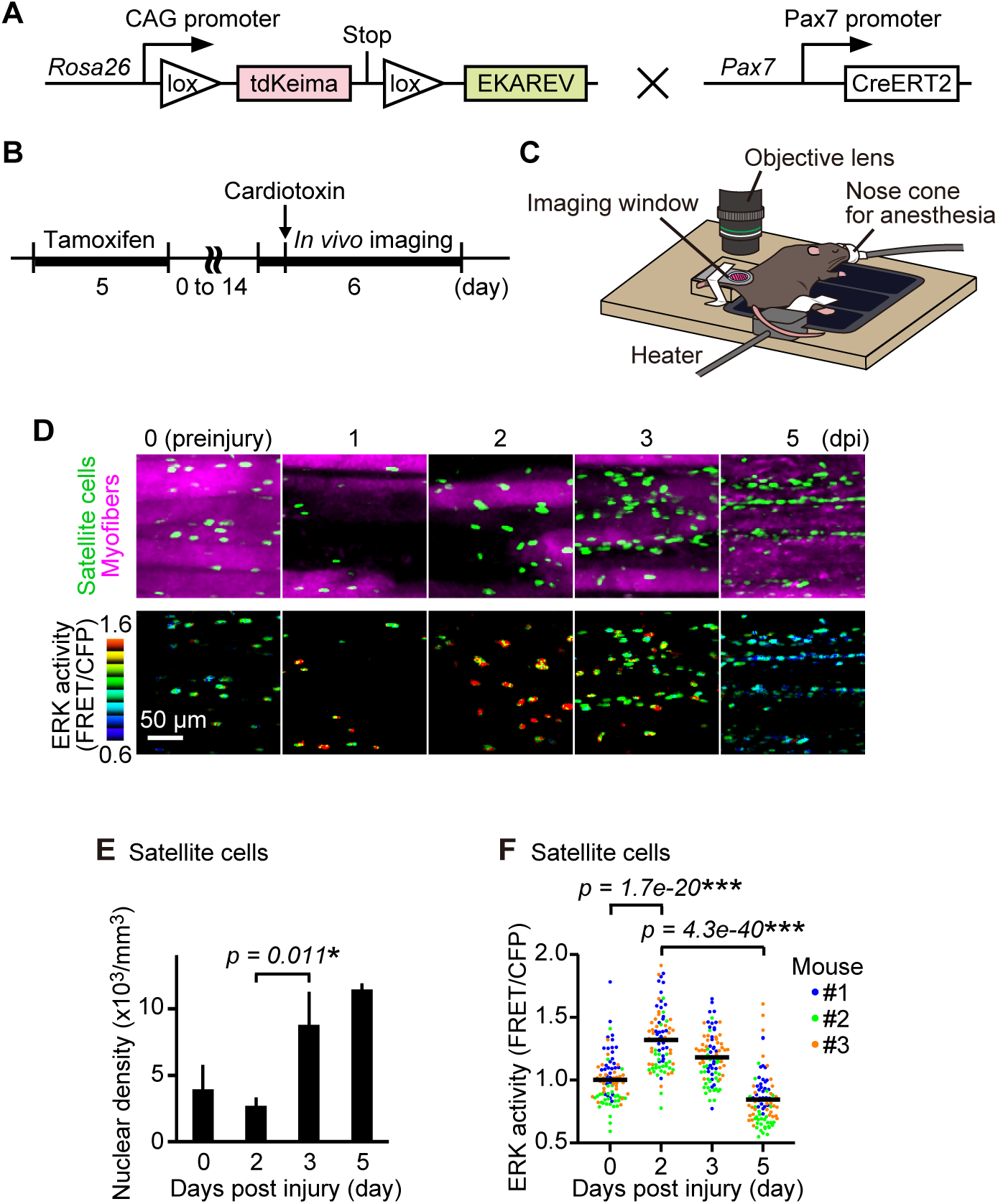
ERK is activated during muscle regeneration. (**A**) Scheme of R26R-EKAREV/Pax7-CreERT2 mice. (**B**) Experimental scheme of Cre-mediated recombination and *in vivo* imaging of skeletal muscle regeneration. (**C**) Layout for the *in vivo* imaging system. The muscle under the imaging window was observed with a two-photon microscope repetitively. (**D**) Representative images of myogenic progenitor cells at 0, 1, 2, 3, and 5 days post injury (dpi). Biceps femoris muscles were imaged as indicated time points, and shown in maximum intensity projection images of 30 μm z-stack with 2 μm intervals. EKAREV-NLS was used to monitor the biosensor in the nucleus. Green and magenta cells in merged images represent myogenic progenitor cells and the myofibers, respectively (top panels). ERK activity (FRET/CFP) images of myogenic progenitor cells shown in the intensity-modulated display (IMD) mode (bottom panels). (**E**) Averaged nuclear density of myogenic progenitor cells calculated from the z-stack images (bars, SDs; N = 3 mice for each day; *p < 0.05; p value is given with an asterisk). (**F**) ERK activity (FRET/CFP) of myogenic progenitor cells. Different color represents datasets from a different mouse (bars, averages; N = 3 mice for each day; ***p < 0.001; p values are given with asterisks).

During muscle regeneration, ERK activity (FRET/CFP) in satellite cells was maximally increased at 2 dpi and decreased below the basal level at 5 dpi (Fig. 1D and 1F). Statistical differences were found among every different pair of days. Collectively, our results indicate that ERK activation precedes proliferation in myogenic satellite cells.

### Immediate ERK activity is required for migration in some satellite cells but not in all satellite cells

Since ERK activity regulates both cell migration and proliferation (Suzuki et al., 2000); (Jones et al., 2001), we first tested the relationship between ERK activity and cell migration speed (Fig. 2A). To examine the migration speed of myogenic progenitor cells, the speed was calculated from the displacement of nuclear centroids tracked more than 1 hour and divided by the time. The migration speed was significantly and maximally increased at 3 dpi and decreased at 5 dpi (Fig. 2B). Because ERK activity was already increased at 2 dpi (Fig. 1F), this observation indicates that ERK activation precedes the increase in migration speed as well as proliferation, in myogenic progenitor cells during muscle regeneration.

**Figure 2.**
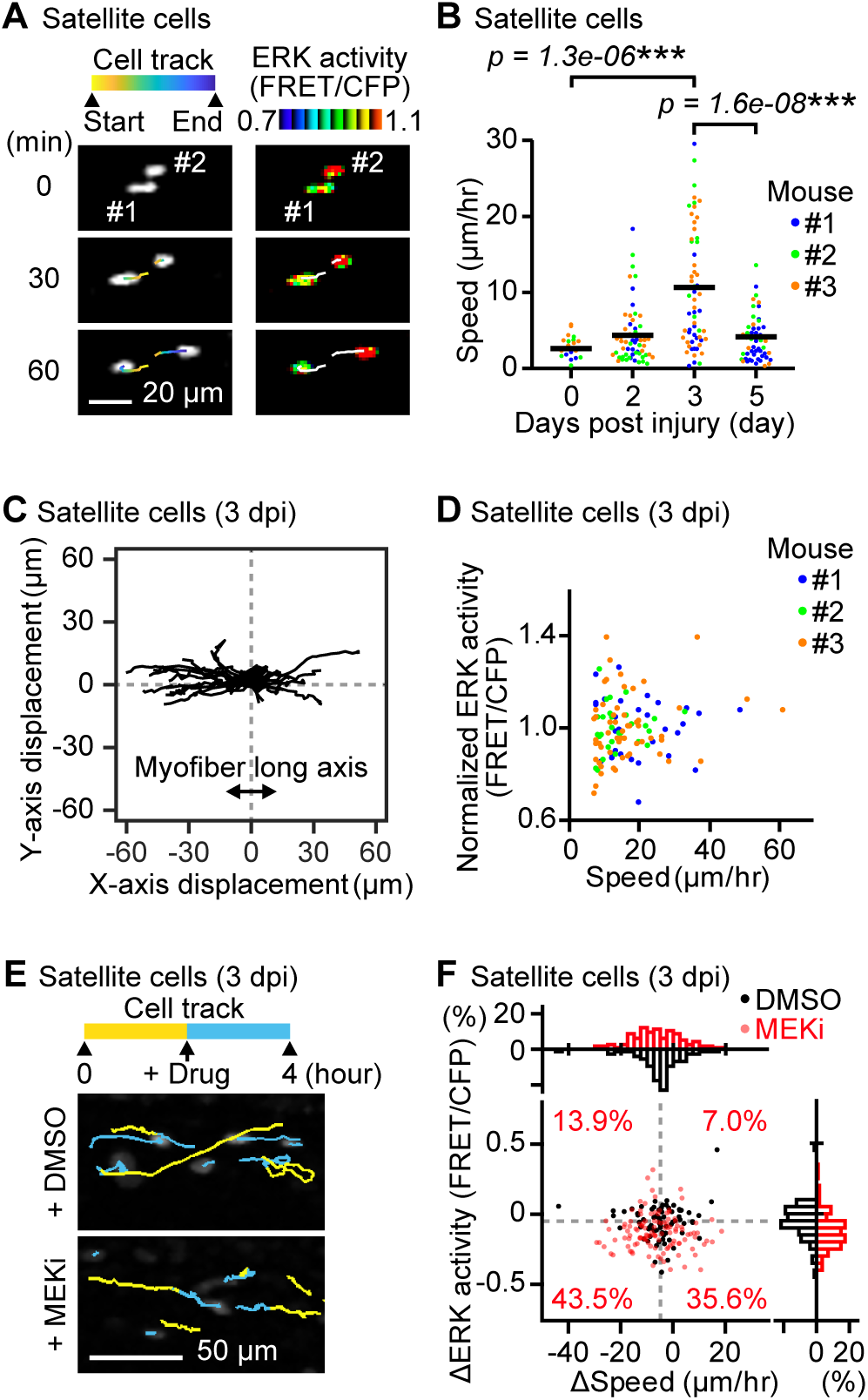
Immediate ERK activity is required for migration in some satellite cells but not in all satellite cells. (**A**) Representative time-lapse images of satellite cells (white dots) and their cell tracks (pseudo-colored lines) (left). FRET/CFP ratio images of satellite cells (IMD mode dots) and their cell tracks (white lines) (right). (**B**) Migration speed of myogenic progenitor cells, which was calculated from the displacement of EKAREV-NLS centroids tracked more than 1 hour and divided by the time. Different color represents datasets from a different mouse (bars, averages; N = 3 mice for each day; ***p < 0.001; p values are given with asterisks). (**C**) Representative cell tracks for 2 hours. X-axis corresponds to the long axis of myofibers. (**D**) Scatter plot of normalized ERK activity (FRET/CFP) against migration speed in migrating satellite cells. Satellite cells with a speed of more than 7 μm/hr were defined as “migrating” and taken into account. ERK activity was normalized by the averaged ERK activity of each mouse. Different color represents datasets from a different mouse (N = 3 mice). (**E**) Representative images of satellite cells (white dots) and their cell tracks (two-colored lines). Yellow lines indicate cell tracks during the first two hours. Blue lines indicate cell tracks during the latter two hours after treatment with DMSO (1 mL/kg) or a MEK inhibitor (PD0325901, 5 mg/kg). (**F**) The difference in migration speed and ERK activity in satellite cells, calculated by subtracting values before MEKi treatment from values after MEKi treatment. Gray dashed lines indicate the median of ERK activity and migration speed in DMSO group. Percentages of each cell groups after MEKi treatment are indicated in the scatter plot. Histograms of the difference in migration speed and ERK activity are shown at the top and right side of the figure, respectively (N = 4 mice for DMSO group; N =3 mice for MEKi group).

Interestingly, we found that satellite cells migrate predominantly along the long axis of myofibers (Fig. 2C), consistent with the finding that extracellular matrix of the basal laminae around myofibers serve as a guide for satellite cells to migrate (Webster et al., 2016). Moreover, the direction of satellite cell migration was not biased toward either of the ends along the long axis of myofibers (Fig. 2C). This result suggests that, at least in muscle regeneration at 3 dpi, satellite cell migration is governed by random walk rather than by chemotaxis.

Although ERK has been reported to promote myoblast migration (Suzuki et al., 2000), to what extent ERK activity is required for satellite cell migration is not completely understood. Therefore, we examined the relationship between ERK activity and speed in satellite cells. Unexpectedly, we failed to observe strong correlation between migration speed and ERK activity at 3 dpi (Fig. 2D, only satellite cells with a speed of more than 7 μm/hr were defined as “migrating” and analyzed). This suggests that migration speed of satellite cells is not immediately determined by ERK activity. We next tested for immediate ERK activity requirement in satellite cell migration by acutely inhibiting MEK, a kinase of ERK, at 3 dpi (Fig. 2E and 2F). A MEK inhibitor treatment only moderately decreased the speed in migrating satellite cells (top histogram, Fig. 2F). Some satellite cells (43.5%) decreased in migration speed and ERK activity (bottom left cell population in scatter plot, Fig. 2F). However, it is important to note that many other migrating satellite cells (35.6%) did not alter their speed after MEK inhibitor treatment, even though ERK activity was significantly decreased (bottom right cell population in scatter plot, Fig. 2F). These results indicate that immediate ERK activity may regulate migration speed in some satellite cells but not in all satellite cells.

### ERK activation is required for the G1/S transition *in vivo*

Due to the lack of correlation in immediate ERK activity and migration speed, we speculated that ERK promotes cell migration through its transcriptional targets (Fig. 2). This is consistent with our observation that that there was a one-day gap between the peak of ERK activity and the peak of cell migration speed (Fig. 1). We thus focused on cell-cycle progression, a key long-term process that is linked to ERK-mediated transcription. First, to clarify the role of ERK in cell-cycle progression *in vivo*, we inhibited ERK activity in R26Fucci2aR/Pax7-CreERT2 mice that expressed a cell cycle indicator, Fucci, in Pax7-expressing satellite cells. Fucci2a is composed of two chimeric proteins, mCherry-hCdt1 and mVenus-hGeminin, which accumulate reciprocally in the nucleus of the cells during the cell cycle, labeling the nuclei of G0/G1 phase cells with mCherry and those of S/G2/M phase cells with mVenus. The proportion of cells expressing mCherry-hCdt1 and cells expressing mVenus-hGeminin was analyzed after ERK activity was suppressed by a MEK inhibitor, PD0325901 (Fig. 3A). Fixed muscle was cleared by CUBIC reagents to obtain the broad cross-sectional area of the tissue. By MEK inhibitor treatment at 2 and 2.5 dpi, the density of cells expressing mVenus-hGeminin was decreased at 3 dpi (Fig. 3B and 3C), suggesting that satellite cells were arrested at the G1/S boundary. This result indicates that ERK activation is required for the G1/S transition *in vivo* during muscle regeneration.

**Figure 3.**
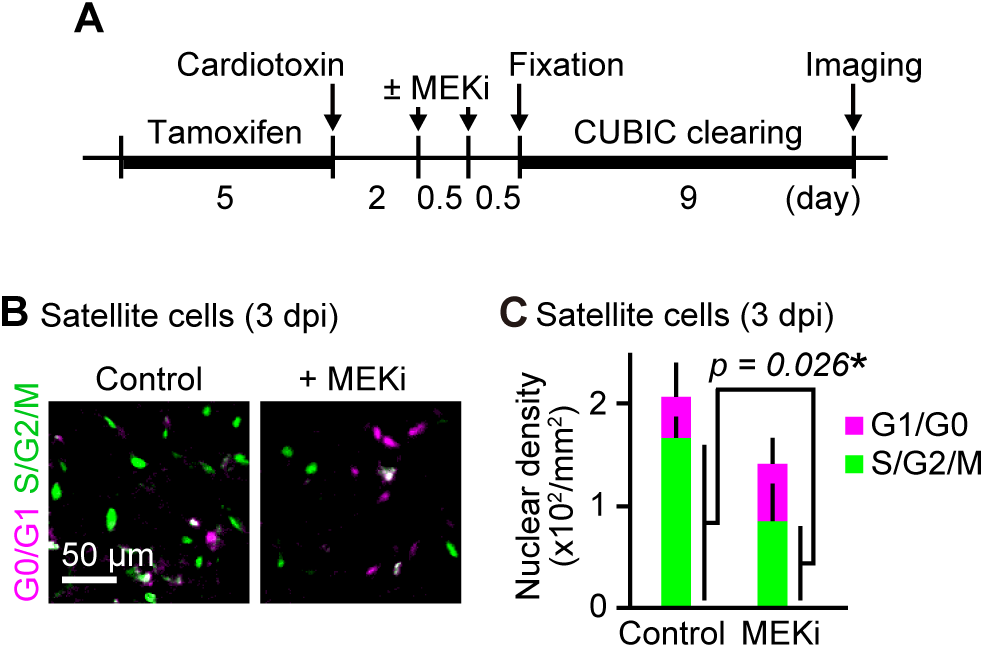
ERK activation is required for the G1/S transition. (**A**) Experimental scheme of Cre-mediated recombination and tissue clearing. Mice were injected with or without a MEK inhibitor (PD0325901, 5 mg/kg) at 2 and 2.5 dpi, and fixed at 3 dpi. (**B** and **C**) Representative images (B) and averaged nuclear density (C) of regenerating regions in the mouse skeletal muscle expressing Fucci in satellite cells. Magenta and green colors represent cells in the G0/G1 and the S/G2/M phase, respectively. Mice were analyzed according to the experimental scheme described in (A) (bars, SDs; N = 3 mice for each group; *p < 0.05; p value is given with an asterisk).

### Migration speed increases in the S/G2 phase

From these data, we hypothesized that ERK promotes cell cycle progression from the G0/G1 to S phase, which precedes the peak of cell migration speed. To further investigate the relationship between cell cycle and migration in satellite cells, progression of cell cycle phase during muscle regeneration was examined using R26Fucci2aR/Pax7-CreERT2 mice. Again, Fucci2aR was expressed in satellite cells by injection of tamoxifen. Then, skeletal muscle damage was induced by cardiotoxin injection. At 0 dpi, almost all of the cells were mCherry positive, i.e., in G0 phase (Fig. 4A and 4B). The cells expressing mVenus-hGeminin increased at 2 to 3 dpi and decreased at 5 dpi. These data indicate satellite cells mainly divide from 1 to 4 dpi.

**Figure 4.**
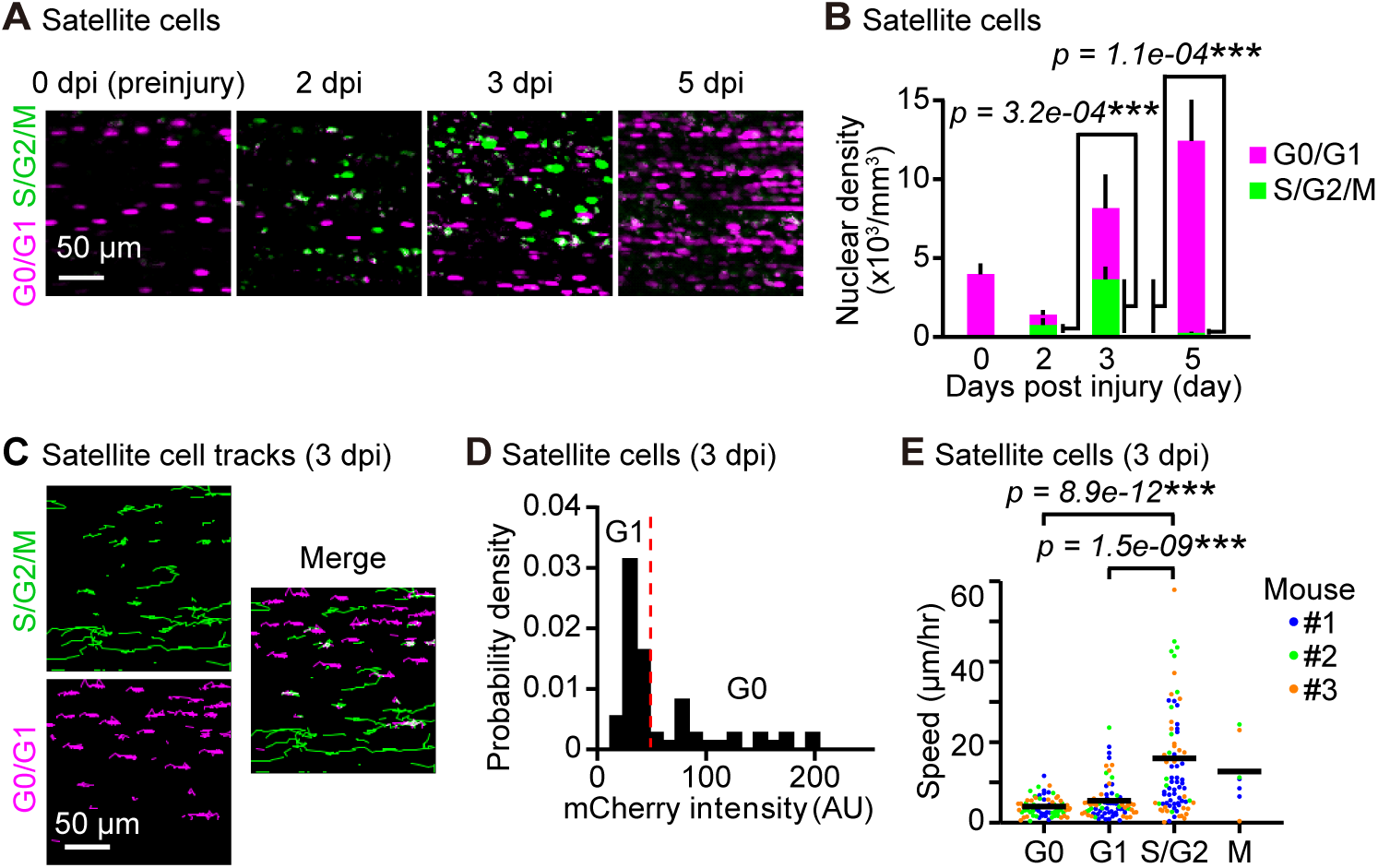
Migration speed of satellite cells increases in the S/G2 phase. (**A**) Representative images of satellite cells expressing Fucci at 0, 2, 3, and 5 dpi. Biceps femoris muscles were imaged as indicated time points, and shown in maximum intensity projection images of 100 μm z-stack with 2 μm intervals. Magenta and green dots indicate cells in the G0/G1 phase and those in the S/G2/M phase, respectively. (**B**) Averaged nuclear densities of satellite cells expressing Fucci calculated from the z-stack images (bars, SDs; N = 3 mice for each day; ***p < 0.001; p value is given with asterisks). (**C**) Representative images of cell trajectories for 4 hours at 3 dpi. Magenta and green lines indicate the trajectories of cells in the G0/G1 phase and those in the S/G2/M phase, respectively. (**D**) Representative probability density distribution of the mCherry-hCdt1 intensity. A red dashed line indicates a threshold to discriminate cells in the G0 and G1 phase. The threshold was defined as an intersection of two Gaussian distributions fitted to the data. (**E**) Migration speed of satellite cells expressing Fucci during each cell cycle phase at 3 dpi. Cells in the G0 and G1 phase were discriminated by the threshold determined in (D). Cells in the M phase was discriminated from cells in the S/G2 phase by cytosolic distribution and subsequent disappearance of mVenus-hGeminin. Different color represents datasets from different mice (bars, averages; N = 3 mice for each day; ***p < 0.001; p values are given with asterisks).

Next, we asked whether the migration speed varies depending on the cell cycle. For this purpose, we focused on 3 dpi because satellite cells are both in G1/S and G2 phases (Fig. 4B). We tracked mCherry-positive or mVenus-positive cells for 4 hours (Fig. 4C). Time-lapse imaging of the skeletal muscle of R26Fucci2aR/Pax7-CreERT2 mice revealed that S/G2/M cells expressing the mVenus-hGeminin migrate faster than G0/G1 cells expressing mCherry-hCdt1. The cells were classified into G0, G1, S/G2 and M phases to further examine the cell cycle dependence of migration speed. Cells that do not express Ki67 (and thus identified as G0 cells) have been previously reported to express higher levels of mKO2-hCdt1 (Tomura et al., 2013). Thus, we classified cells expressing higher and lower levels of mCherry-hCdt1 as cells in G0 and G1 phase, respectively (Fig. 4D). Cells in the M phase were discriminated from cells in the S/G2 phase by nuclear membrane breakdown and subsequent disappearance of mVenus-hGeminin. With these analyses, we found that the migration speed maximally increased in the S/G2 phase and decreased in the M phase and G1 phase, and reached the minimum in the G0 phase (Fig. 4E).

### CDK2 promotes satellite cell migration during muscle regeneration

These results motivated us to search for a mechanism underlying cell cycle-dependent migration. To this end, we examined the contribution of CDKs, whose activities are tightly controlled throughout the cell cycle (Malumbres and Barbacid, 2009). To test the hypothesis that a downstream substrate of CDK directly regulates cell migration, CDK inhibitors were injected in mice during *in vivo* imaging at 3 dpi. The difference in migration speed of each cell was plotted against the difference in ERK activity (Fig. 5A, 5B, and 5C). To clarify the effects of CDK inhibitors on cell migration, we focused on the migrating satellite cells that decreased their speed more than 7 μm/hr after the inhibitor treatment, and defined as “decelerated”. Gray dashed lines denote the threshold for classifying “decelerated” population (Fig. 5A, 5B, and 5C). Interestingly, decelerated cell population was increased by a CDK1/2 inhibitor, roscovitine, but not by a CDK1 inhibitor or a CDK4/6 inhibitor (Fig. 5D). From these results, we speculated that CDK2 could promote satellite cell migration. This hypothesis is also advocated by the facts that CDK1 is most activated in M phase and that migration speed is higher in the S/G2 than in M phase (Fig. 4E). Given that roscovitine is a kinase inhibitor, this result implies that phosphorylation of a CDK2 downstream substrate promotes satellite cell migration in the S/G2 phase during muscle regeneration.

**Figure 5.**
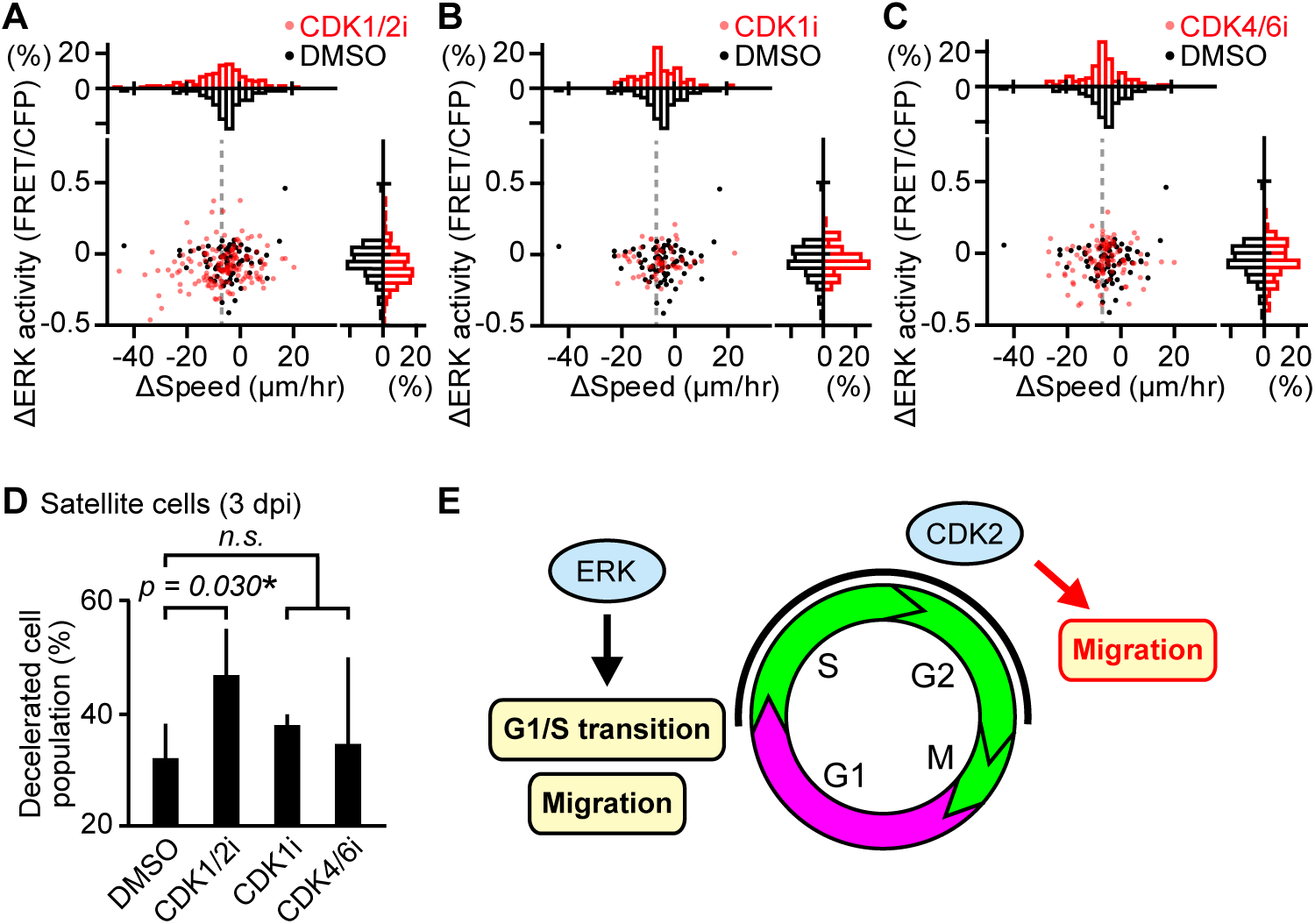
CDK2 promotes cell migration during muscle regeneration. (**A, B**, and **C**) The difference in migration speed and ERK activity in satellite cells, calculated by subtracting values before the drug treatment from values after the drug treatment. Gray dashed lines indicate 7 μm/hr of decrease in migration speed. Histograms of the difference in migration speed and ERK activity are shown at the top and right side of the figure, respectively (N = 4 mice for DMSO group; N = 4 mice for CDK1/2i group; N =3 mice for CDK1i group; N = 3 mice for CDK4/6i group). Mice expressing EKAREV-NLS in satellite cells were injected intravenously with DMSO (1 mL/kg), CDK1/2 inhibitor (roscovitine, 0.05 mg/kg), CDK1 inhibitor (RO-3306, 1 mg/kg), or CDK4/6 inhibitor (palbociclib, 1 mg/kg) during *in vivo* imaging at 3 dpi. (**D**) Percentage of decelerated cell population after DMSO or CDK inhibitors. Migrating satellite cells that decreased their speed more than 7 μm/hr are defined as “decelerated” and taken into account (bars, SDs; *p < 0.05; n.s., not significant; p value is given with an asterisk). (**E**) Schematic model of cell cycle progression and cell migration by ERK and CDK2 in satellite cells during muscle regeneration.

## Discussion

Based on our results, we propose two modes of ERK action in satellite cell during regeneration, in the early and later phase. The effect of ERK in the early phase coincides with satellite cell activation and promotes cell migration, whereas the effect of ERK the later phase promotes the G1/S transition and cell migration though CDK2 activation (Fig. 5E).

We demonstrate, for the first time, that ERK is activated upon satellite cell activation during muscle regeneration *in vivo*. Our data suggests that ERK activation precedes other regulators of muscle regeneration. ERK activation peaks at 2 dpi, while the myogenic master transcription factor MyoD expression peaks at 3 to 4 dpi (Ogawa et al., 2015) and other mitogen-activated protein kinase p38 peaks at 7 to 14 dpi (Ruiz-Bonilla et al., 2008). This supports the notion that ERK is activated early in muscle regeneration, when satellite cells exit from quiescence in response to injury.

The varied effects of ERK on cell migration among satellite cells *in vivo* could be caused by the difference in focal adhesion signaling. Pro-migratory functions of ERK and the responsible substrates have been characterized in numerous cell types. Among the identified substrates, two focal adhesion–associated proteins, FAK and paxillin are most likely to be involved in ERK-induced cell migration in satellite cells. ERK is suggested to interact with FAK/paxillin and promote cell migration by enhancing focal adhesion turnover and membrane protrusion at the front of the cells (Hauck et al., 2000); (Liu et al., 2002); (Subauste et al., 2004); (Teranishi et al., 2009); (Singh et al., 2019). Functional effects of FAK/paxillin were corroborated by *in vivo* studies showing that targeted deletion of FAK in satellite cells impairs skeletal muscle regeneration (Quach et al., 2009), and that paxillin is hyper-phosphorylared in dystrophin-deficient *mdx* muscle (Sen et al., 2011). Therefore, the difference in abundance of FAK and paxillin could explain the difference in the ERK contribution on cell migration among satellite cells.

Multiple lines of evidence support a pro-migratory role of CDK2 though stathmin, a phosphorylation-regulated tubulin-binding protein. First of all, stathmin is suggested to be phosphorylated at Ser25 by CDK2, in a consensus CDK/MAPK phosphorylation motif, PXS*P (Chi et al., 2008). In agreement with our model, several papers have demonstrated that p27, the cyclin-dependent kinase inhibitor, inhibits cell migration though CDK2 and stathmin (Baldassarre et al., 2005); (Schiappacassi et al., 2008); (Schiappacassi et al., 2011); (Nadeem et al., 2013). Furthermore, p27 knockout mouse showed increased body weight along with muscle weight (Kiyokawa et al., 1996), whereas stathmin knockout mouse developed age-dependent myopathy (Liedtke et al., 2002). More interestingly, the expression of stathmin has been suggested to increase as myoblasts undergo differentiation (Balogh et al., 1996); (Gonnet et al., 2008); (Casadei et al., 2009). Further study is needed regarding the mechanism by which CDK2 promote cell migration.

We speculate that the cell cycle-dependent migration in satellite cells may contribute to efficient regeneration and differentiation, mediated by CDK2, p21, and the myogenic master transcription factor MyoD. Of note, crosstalk between cell cycle regulators and myogenic regulatory factors has been well characterized *in vitro*. Expression of MyoD peaks in mid-G1, and is reduced to its minimum level at the G1/S transition (Kitzmann et al., 1998). In late G1, MyoD is degraded by the ubiquitin proteasome system, which is triggered by cyclin E/CDK2-dependent phosphorylation of MyoD at Ser200 (Song et al., 1998); (Kitzmann et al., 1999); (Tintignac et al., 2004). In turn, MyoD inhibits CDK2 activity by inducing expression of the cyclin-dependent kinase inhibitor p21 (Halevy et al., 1995); (Guo et al., 2015). Although satellite cells need to proliferate and migrate into the site of injury, they also need to stop migrating and differentiate into myotubes, by fusing to each other or to the remaining myofibers. We speculated that such migration control is important especially where cells migrate stochastically along the long axis of myofibers (Fig. 2C). Thus, higher motility of satellite cells in S/G2 would help to supply satellite cells at the site of injury, whereas lower motility of satellite cells in G1 would be beneficial to induce efficient differentiation into myofibers.

In summary, we demonstrated that satellite cells migrate in a cell cycle-dependent manner and that both ERK and CDK2 contribute to promoting their migration during muscle regeneration *in vivo*, which may provide the mechanism underlying efficient muscle regeneration. These findings highlight the importance of studying molecular activity, cell migration, and cell-cycle phases in living tissue with intravital imaging.

## Materials and Methods

### Reagents

PD0325901 (FUJIFILM Wako Pure Chemical Corporation, Osaka, Japan), roscovitine (Sigma-Aldrich, St. Louis, MO), RO-3306 (Tokyo Chemical Industry, Tokyo, Japan), and palbociclib (Chemietek, Indianapolis, IN) were applied as inhibitors for MEK, CDK1/2, CDK1, and CDK4/6, respectively.

### Transgenic mice

Gt(ROSA)26Sor^tm1(CAG-loxP-tdKeima-loxP-EKAREV-NLS)^ (hereinafter called R26R-EKAREV-NLS) mice have been developed previously (Konishi et al., 2018). These mouse lines are designed to express the tdKeima fluorescent protein before Cre-mediated recombination and EKAREV after recombination, under the CAG promoter in the *ROSA26* locus. Gt(ROSA)26Sor^tm1(Fucci2aR)Jkn^ (hereinafter called R26Fucci2aR) mice have been developed previously (Mort et al., 2014). B6;129-Pax7^tm2.1(cre/ERT2)Fan^/J (hereinafter called Pax7-CreERT2) mice have been developed previously (Lepper et al., 2009) and were provided by Atsuko Sehara-Fujisawa (Kyoto University, Kyoto, Japan). This mouse line is designed to express a tamoxifen-inducible Cre recombinase–oestrogen receptor fusion protein, CreERT2 under the endogenous promoter in the *Pax7* locus.

To develop transgenic mice expressing EKAREV-NLS or Fucci in satellite cells specifically, R26R-EKAREV-NLS or R26Fucci2aR mice were crossed with Pax7-CreERT2 mice. To induce Cre mediated recombination, tamoxifen (Sigma-Aldrich) dissolved in corn oil (Sigma-Aldrich) were injected into intraperitoneally (75 mg/kg) once a day consecutively for five days. Mice were housed in a specific-pathogen-free facility and received a routine chow diet and water *ad libitum*. Adult female and male mice of 2 to 6 months of age were used for the *in vivo* imaging. The animal protocols were reviewed and approved by the Animal Care and Use Committee of Kyoto University Graduate School of Medicine (No.14079, 15064, 16038, 17539, and 18086).

### Muscle injury with cardiotoxin

To investigate the muscle regeneration, muscle damage was induced by cardiotoxin. The skin over the skeletal muscle was shaved and cleaned with 70% ethanol. The skeletal muscle was injected with 10 μL of cardiotoxin (Sigma-Aldrich or Latoxan, Portes lès Valence, France) in DDW (1 mg/mL).

### *In vivo* imaging of skeletal muscle

For repetitive observations, the custom-made imaging window were implanted in the femoral region as described previously (Takaoka et al., 2016) before cardiotoxin injection. For a single observation, the skin over the tibialis anterior (TA) was shaved and incised to expose approximately 1 cm^2^ of the TA muscle as described previously (Konagaya et al., 2017) after cardiotoxin injection. Mice were anaesthetized with 1 to 1.5% isoflurane (FUJIFILM Wako Pure Chemical Corporation) mixed with oxygen delivered at 1 L/min. Drugs were injected intravenously during imaging.

### Tissue clearing

For tissue clearing, the TA muscle was collected from mice and fixed in 4% PFA overnight in 4°C. The fixed organs were immersed in CUBIC-1 reagent for 5 days and then further immersed in CUBIC-2 reagent. ScaleCUBIC-1 (reagent-1A) was prepared as a mixture of 10 wt% urea (Nacalai Tesque, Kyoto, Japan), 5 wt% N, N, N’, N’-tetrakis (2-hydroxypropyl) ethylenediamine (Tokyo Chemical Industry), 10 wt% Triton X-100 (Nacalai Tesque), and 25 mM NaCl (Nacalai Tesque). ScaleCUBIC-2 (reagent 2) was prepared as a mixture of 50 wt% sucrose (Nacalai Tesque), 25 wt% urea, 10 wt% 2, 2’, 2”-nitrilotriethanol (FUJIFILM Wako Pure Chemical Corporation), and 0.1% (v/v) Triton X-100 (Susaki et al., 2014).

### Two-photon excitation microscopy

For repetitive observations, living mice were observed with an FV1200MPE-BX61WI upright microscope (Olympus, Tokyo, Japan) equipped with an XLPLN25XWMP water-immersion objective lens (Olympus), where the pixel size was 1.59 um/pixel. For a single observation, living mice were observed with an FV1200MPE-IX83 inverted microscope (Olympus) equipped with a UPlanSApo 30x/1.05NA silicon oil-immersion objective lens (Olympus), where the pixel size was 1.325 um/pixel. The microscopes were equipped with an InSight DeepSee Ultrafast laser (0.95 W at 900 nm) (Spectra Physics, Mountain View, CA). The scan speed was set at 2 to 10 µs/pixel. The excitation wavelength for CFP, GFP, and RFP was 840, 960, and 1040 nm, respectively.

Fluorescent images were acquired with the following filters and mirrors: (1) an infrared-cut filter BA685RIF-3 (Olympus), (2) two dichroic mirrors DM505 (Olympus) and DM570 (Olympus), and (3) four emission filters FF01-425/30 for second harmonic generation (SHG) (Semrock, Rochester, NY), BA460-500 for CFP/SHG (Olympus), BA520-560 for FRET/GFP (Olympus), and 645/60 for RFP (Chroma Technology, Bellows Falls, VT). The microscopes were equipped with a two-channel GaAsP detector unit and two multialkali detectors. FLUOVIEW software (Olympus) was used to control the microscope and to acquire images, which were saved in the multilayer 12-bit tagged image file format.

### Lightsheet microscopy

Images of cleared tissues were acquired with a Lightsheet Z.1 microscope (Zeiss, Oberkochen, Germany) equipped with a single side light sheet and two lenses: an EC Plan-Neofluar 5x/0.16 detection objective lens and LSFM clearing 5x/0.1 illumination objective lens. The excitation wavelength for mVenus and mCherry was 488 and 561 nm, respectively. The light sheet thickness was 12.67 μm. A laser blocking filter, LBF 405/488/561/640, secondary beam splitters, SBS LP490 and SBS LP560, and emission filters, BP505-545 and BP575-615, were used. Images were saved in the multilayer 16-bit tagged image file format. ZEN software (Zeiss) was used to control the microscope and to acquire images. Samples were immersed in a 1:1 mixture of silicon oil TSF4300 (Momentive Performance Materials Japan, Tokyo) and mineral oil (Sigma-Aldrich) during image acquisition.

### Image processing

Acquired images were processed with ImageJ (National Institutes of Health, Bethesda, MD, USA) and MATLAB software (MathWorks, Natick, MA).

ImageJ software was used to obtain x- and y-coordinates of the nuclei centroid. First, z-stack images were aligned using an ImageJ plug-in “Correct 3D drift” (Parslow et al., 2014). The CFP or SHG images were used as landmarks for the correction. Corrected z-stack images were processed with a median filter (5×5×5 pixels) and subtracted background noise with a top-hat filter (11×11 pixels). Filtered images were maximum intensity projected along the z axis. The nuclei were tracked with an ImageJ plug-in “Trackmate” (Tinevez et al., 2017). For efficient tracking, CFP images were contrast adjusted using an ImageJ plug-in “Stack Contrast Adjustment” (Čapek et al., 2006). The parameters in Trackmate were set as follows:

Detector: LoG detector

Estimated blob diameter: 5pixels

Intensity threshold: 0

Median filter: false

Sub-pixel localization: true

Local maxima: 3 (for SECFP in EKAREV-NLS), 5 (for mVenus in Fucci), and 1 (for mCherry in Fucci)

Tracker: Simple LAP tracker

Linking max distance: 10 pixels

Gap-closing max distance: 10 pixels

Gap-closing max frame: 2

MATLAB standard and custom-written scripts were used to obtain the FRET/CFP ratio and the speed. The FRET/CFP ratio was calculated by dividing the averaged FRET intensity by the averaged CFP in the radial distance of 1-pixel from the centroid. The speed was calculated by dividing the displacement of the centroid by the time.

### Statistical analysis

Graphing and statistical analysis was performed with MATLAB software. Statistical differences between two experimental groups were assessed by Student’s two-sample two-sided t-test. Statistical differences among experimental groups more than two were assessed by Scheffe’s F-test. Statistical significances were indicated by asterisks (*p < 0.05; **p < 0.01; ***p < 0.001).

## Acknowledgements

Financial support for this article was provided in the form of JSPS KAKENHI18K07066 (K.T.), JSPS KAKENHI19H00993, JSPS KAKENHI15H05949 “Resonance Bio,” JSPS KAKENHI16H06280 “ABiS,” CRESTJPMJCR1654 (M.M.) grants, and the Fellowship from Astellas Foundation (Y.K.).

## References

Almada, A.E., and Wagers, A.J. (2016). Molecular circuitry of stem cell fate in skeletal muscle regeneration, ageing and disease. Nature Reviews Molecular Cell Biology 17, 267–279.

Baldassarre, G., Belletti, B., Nicoloso, M.S., Schiappacassi, M., Vecchione, A., Spessotto, P., Morrione, A., Canzonieri, V., and Colombatti, A. (2005). p27 Kip1 -stathmin interaction influences sarcoma cell migration and invasion. Cancer Cell 7, 51–63.

Balogh, A., Mège, R.M., and Sobel, A. (1996). Growth and cell density-dependent expression of stathmin in C2 myoblasts in culture. Experimental Cell Research 224, 8–15.

Blau, H.M., Cosgrove, B.D., and Ho, A.T.V. (2015). The central role of muscle stem cells in regenerative failure with aging. Nature Medicine 21, 854–862.

Bouchard, G., Bouvette, G., Therriault, H., Bujold, R., Saucier, C., and Paquette, B. (2013). Pre-irradiation of mouse mammary gland stimulates cancer cell migration and development of lung metastases. British Journal of Cancer 109, 1829–1838.

Burstyn-Cohen, T., and Kalcheim, C. (2002). Association between the cell cycle and neural crest delamination through specific regulation of G1/S transition. Developmental cell 3, 383–395.

Capek, M., Janáček, J., and Kubínová, L. (2006). Methods for compensation of the light attenuation with depth of images captured by a confocal microscope. Microscopy Research and Technique 69, 624–635.

Casadei, L., Vallorani, L., Gioacchini, A.M., Guescini, M., Burattini, S., D’Emilio, A., Biagiotti, L., Falcieri, E., and Stocchi, V. (2009). Proteomics-based investigation in C2C12 myoblast differentiation. European journal of histochemistry: EJH 53, 261–268.

Ceafalan, L.C., Popescu, B.O., and Hinescu, M.E. (2014). Cellular players in skeletal muscle regeneration. Biomed Res Int 2014, 957014.

Chi, Y., Welcker, M., Hizli, A.A., Posakony, J.J., Aebersold, R., and Clurman, B.E. (2008). Identification of CDK2 substrates in human cell lysates. Genome Biology 9, 1–12.

Corcoran, A., Del Maestro, R.F., Berger, M.S., Canoll, P.D., and Bruce, J.N. (2003). Testing the “Go or Grow” hypothesis in human medulloblastoma cell lines in two and three dimension. Neurosurgery 53, 174–185.

Doukas, J., Blease, K., Craig, D., Ma, C., Chandler, L.A., Sosnowski, B.A., and Pierce, G.F. (2002). Delivery of FGF genes to wound repair cells enhances arteriogenesis and myogenesis in skeletal muscle. Molecular Therapy 5, 517–527.

Floss, T., Arnold, H.H., and Braun, T. (1997). A role for FGF-6 in skeletal muscle regeneration. Genes and Development 11, 2040–2051.

Garay, T., Juhász, É., Molnár, E., Eisenbauer, M., Czirók, A., Dekan, B., László, V., Hoda, M.A., Döme, B., Tímár, J., et al. (2013). Cell migration or cytokinesis and proliferation? - Revisiting the “go or grow” hypothesis in cancer cells in vitro. Experimental Cell Research 319, 3094–3103.

Giese, A., Loo, M.A., Tran, N., Haskett, D., Coons, S.W., and Berens, M.E. (1996). Dichotomy of astrocytoma migration and proliferation. International Journal of Cancer 67, 275–282.

Gonnet, F., Bouazza, B., Millot, G.A., Ziaei, S., Garcia, L., Butler-Browne, G.S., Mouly, V., Tortajada, J., Danos, O., and Svinartchouk, F. (2008). Proteome analysis of differentiating human myoblasts by dialysis-assisted two-dimensional gel electrophoresis (DAGE). Proteomics 8, 264–278.

Guo, K., Wang, J., Andrés, V., Smith, R.C., and Walsh, K. (2015). MyoD-induced expression of p21 inhibits cyclin-dependent kinase activity upon myocyte terminal differentiation. Molecular and Cellular Biology 15, 3823–3829.

Haass, N.K., Beaumont, K.A., Hill, D.S., Anfosso, A., Mrass, P., Munoz, M.A., Kinjyo, I., and Weninger, W. (2014). Real-time cell cycle imaging during melanoma growth, invasion, and drug response. Pigment Cell and Melanoma Research 27, 764–776.

Halevy, O., Novitch, B., Spicer, D., Skapek, S., Rhee, J., Hannon, G., Beach, D., and Lassar, A. (1995). Correlation of terminal cell cycle arrest of skeletal muscle with induction of p21 by MyoD. Science 267, 1018–1021.

Hauck, C.R., Hsia, D.A., and Schlaepfer, D.D. (2000). Focal adhesion kinase facilitates platelet-derived growth factor-BB-stimulated ERK2 activation required for chemotaxis migration of vascular smooth muscle cells. Journal of Biological Chemistry 275, 41092–41099.

Hotta, K., Behnke, B.J., Masamoto, K., Shimotsu, R., Onodera, N., Yamaguchi, A., Poole, D.C., and Kano, Y. (2018). Microvascular permeability of skeletal muscle after eccentric contraction-induced muscle injury: in vivo imaging using two-photon laser scanning microscopy. Journal of Applied Physiology 125, 369–380.

Jones, N.C., Fedorov, Y.V., Rosenthal, R.S., and Olwin, B.B. (2001). ERK1/2 is required for myoblast proliferation but is dispensable for muscle gene expression and cell fusion. Journal of Cellular Physiology 186, 104–115.

Kagawa, Y., Matsumoto, S., Kamioka, Y., Mimori, K., Naito, Y., Ishii, T., Okuzaki, D., Nishida, N., Maeda, S., Naito, A., et al. (2013). Cell cycle-dependent Rho GTPase activity dynamically regulates cancer cell motility and invasion in vivo. PLoS ONE 8.

Kitzmann, M., Carnac, G., Vandromme, M., Primig, M., Lamb, N.J.C., and Fernandez, A. (1998). The muscle regulatory factors MyoD and Myf-5 undergo distinct cell cycle-specific expression in muscle cells. Journal of Cell Biology 142, 1447–1459.

Kitzmann, M., Vandromme, M., Schaeffer, V., Carnac, G., Labbé, J.C., Lamb, N., and Fernandez, A. (1999). cdk1- and cdk2-mediated phosphorylation of MyoD Ser200 in growing C2 myoblasts: role in modulating MyoD half-life and myogenic activity. Molecular and cellular biology 19, 3167–3176.

Kiyokawa, H., Kineman, R.D., Manova-Todorova, K.O., Soares, V.C., Huffman, E.S., Ono, M., Khanam, D., Hayday, A.C., Frohman, L.A., and Koff, A. (1996). Enhanced growth of mice lacking the cyclin-dependent kinase inhibitor function of P27Kip1. Cell 85, 721–732.

Konagaya, Y., Terai, K., Hirao, Y., Takakura, K., Imajo, M., Kamioka, Y., Sasaoka, N., Kakizuka, A., Sumiyama, K., Asano, T., et al. (2017). A Highly Sensitive FRET Biosensor for AMPK Exhibits Heterogeneous AMPK Responses among Cells and Organs. Cell Reports 21, 2628–2638.

Konishi, Y., Terai, K., Furuta, Y., Kiyonari, H., Abe, T., Ueda, Y., Kinashi, T., Hamazaki, Y., Takaori-Kondo, A., and Matsuda, M. (2018). Live-Cell FRET Imaging Reveals a Role of Extracellular Signal-Regulated Kinase Activity Dynamics in Thymocyte Motility. Food Science and Human Wellness 10, 98–113.

Koyama, T., Nakaoka, Y., Fujio, Y., Hirota, H., Nishida, K., Sugiyama, S., Okamoto, K., Yamauchi-Takihara, K., Yoshimura, M., Mochizuki, S., et al. (2008). Interaction of scaffolding adaptor protein Gab1 with tyrosine phosphatase SHP2 negatively regulates IGF-I-dependent myogenic differentiation via the ERK1/2 signaling pathway. Journal of Biological Chemistry 283, 24234–24244.

Kuang, S., Chargé, S.B., Seale, P., Huh, M., and Rudnicki, M.A. (2006). Distinct roles for Pax7 and Pax3 in adult regenerative myogenesis. Journal of Cell Biology 172, 103–113.

Lau, J., Goh, C.C., Devi, S., Keeble, J., See, P., Ginhoux, F., and Ng, L.G. (2016). Intravital multiphoton imaging of mouse tibialis anterior muscle. IntraVital 5, e1156272–e1156272.

Le Grand, F., Grifone, R., Mourikis, P., Houbron, C., Gigaud, C., Pujol, J., Maillet, M., Pagès, G., Rudnicki, M., Tajbakhsh, S., et al. (2012). Six1 regulates stem cell repair potential and self-renewal during skeletal muscle regeneration. Journal of Cell Biology 198, 815–832.

Lepper, C., Conway, S.J., and Fan, C.-M. (2009). Adult satellite cells and embryonic muscle progenitors have distinct genetic requirements. Nature 460, 627–631.

Liedtke, W., Leman, E.E., Fyffe, R.E.W., Raine, C.S., and Schubart, U.K. (2002). Stathmin-deficient mice develop an age-dependent axonopathy of the central and peripheral nervous systems. American Journal of Pathology 160, 469–480.

Liu, Z.X., Yu, C.F., Nickel, C., Thomas, S., and Cantley, L.G. (2002). Hepatocyte growth factor induces ERK-dependent paxillin phosphorylation and regulates paxillin-focal adhesion kinase association. Journal of Biological Chemistry.

Malumbres, M., and Barbacid, M. (2009). Cell cycle, CDKs and cancer: A changing paradigm. Nature Reviews Cancer 9, 153–166.

Michailovici, I., Harrington, H.A., Azogui, H.H., Yahalom-Ronen, Y., Plotnikov, A., Ching, S., Stumpf, M.P.H., Klein, O.D., Seger, R., and Tzahor, E. (2014). Nuclear to cytoplasmic shuttling of ERK promotes differentiation of muscle stem/progenitor cells. Development 141, 2611–2620.

Mort, R.L., Ford, M.J., Sakaue-Sawano, A., Lindstrom, N.O., Casadio, A., Douglas, A.T., Keighren, M.A., Hohenstein, P., Miyawaki, A., and Jackson, I.J. (2014). Fucci2a: A bicistronic cell cycle reporter that allows Cre mediated tissue specific expression in mice. Cell Cycle 13, 2681–2696.

Nadeem, L., Brkic, J., Chen, Y.F., Bui, T., Munir, S., and Peng, C. (2013). Cytoplasmic mislocalization of p27 and CDK2 mediates the anti-migratory and anti-proliferative effects of Nodal in human trophoblast cells. Journal of Cell Science 126, 445–453.

Neuhaus, P., Oustanina, S., Loch, T., Kruger, M., Bober, E., Dono, R., Zeller, R., and Braun, T. (2003). Reduced Mobility of Fibroblast Growth Factor (FGF)-Deficient Myoblasts Might Contribute to Dystrophic Changes in the Musculature of FGF2/FGF6/mdx Triple-Mutant Mice. Molecular and Cellular Biology 23, 6037–6048.

Nobis, M., Warren, S.C., Lucas, M.C., Murphy, K.J., Herrmann, D., and Timpson, P. (2018). Molecular mobility and activity in an intravital imaging setting – implications for cancer progression and targeting. Journal of Cell Science 131, jcs206995–jcs206995.

Ogawa, R., Ma, Y., Yamaguchi, M., Ito, T., Watanabe, Y., Ohtani, T., Murakami, S., Uchida, S., De Gaspari, P., Uezumi, A., et al. (2015). Doublecortin marks a new population of transiently amplifying muscle progenitor cells and is required for myofiber maturation during skeletal muscle regeneration. Development (Cambridge) 142, 51–61.

Oustanina, S., Hause, G., and Braun, T. (2004). Pax7 directs postnatal renewal and propagation of myogenic satellite cells but not their specification. EMBO Journal 23, 3430–3439.

Parslow, A., Cardona, A., and Bryson-Richardson, R.J. (2014). Sample Drift Correction Following 4D Confocal Time-lapse Imaging. Journal of Visualized Experiments, 1–4.

Pittet, M.J., and Weissleder, R. (2011). Intravital imaging. Cell 147, 983–991.

Quach, N.L., Biressi, S., Reichardt, L.F., Keller, C., and Rando, T.A. (2009). Focal adhesion kinase signaling regulates the expression of caveolin 3 and beta1 integrin, genes essential for normal myoblast fusion. Molecular biology of the cell 20, 3422–3435.

Rajan, S.G., Gallik, K.L., Monaghan, J.R., Uribe, R.A., Bronner, M.E., and Saxena, A. (2018). Tracking neural crest cell cycle progression in vivo. Genesis 56.

Rommel, C., Clarke, B.A., Zimmermann, S., Nuñez, L., Rossman, R., Reid, K., Moelling, K., Yancopoulos, G.D., and Glass, D.J. (1999). Differentiation stage-specific inhibition of the Raf-MEK-ERK pathway by Akt. Science 286, 1738–1741.

Ruiz-Bonilla, V., Perdiguero, E., Gresh, L., Serrano, A.L., Zamora, M., Sousa-Victor, P., Jardí, M., Wagner, E.F., and Muñoz-Cánoves, P. (2008). Efficient adult skeletal muscle regeneration in mice deficient in p38β, p38γ and p38δ MAP kinases. Cell Cycle 7, 2208–2214.

Sakaue-Sawano, A., Kurokawa, H., Morimura, T., Hanyu, A., Hama, H., Osawa, H., Kashiwagi, S., Fukami, K., Miyata, T., Miyoshi, H., et al. (2008). Visualizing spatiotemporal dynamics of multicellular cell-cycle progression. Cell 132, 487–498.

Schiappacassi, M., Lovat, F., Canzonieri, V., Belletti, B., Berton, S., Di Stefano, D., Vecchione, A., Colombatti, A., and Baldassarre, G. (2008). p27Kip1 expression inhibits glioblastoma growth, invasion, and tumor-induced neoangiogenesis. Molecular Cancer Therapeutics 7, 1164–1175.

Schiappacassi, M., Lovisa, S., Lovat, F., Fabris, L., Colombatti, A., Belletti, B., and Baldassarre, G. (2011). Role of T198 modification in the regulation of p27Kip1 protein stability and function. PLoS ONE 6.

Seale, P., Sabourin, L.A., Girgis-Gabardo, A., Mansouri, A., Gruss, P., and Rudnicki, M.A. (2000). Pax7 is required for the specification of myogenic satellite cells. Cell 102, 777–786.

Sen, S., Tewari, M., Zajac, A., Barton, E., Sweeney, H.L., and Discher, D.E. (2011). Upregulation of paxillin and focal adhesion signaling follows Dystroglycan Complex deletions and promotes a hypertensive state of differentiation. European Journal of Cell Biology 90, 249–260.

Singh, J., Sharma, K., Frost, E.E., and Pillai, P.P. (2019). Role of PDGF-A-Activated ERK Signaling Mediated FAK-Paxillin Interaction in Oligodendrocyte Progenitor Cell Migration. Journal of molecular neuroscience: MN 67, 564–573.

Song, A., Wang, Q., Goebl, M.G., and Harrington, M.A. (1998). Phosphorylation of Nuclear MyoD Is Required for Its Rapid Degradation. Molecular and Cellular Biology 18, 4994–4999.

Sousa-Victor, P., García-Prat, L., Serrano, A.L., Perdiguero, E., and Muñoz-Cánoves, P. (2015). Muscle stem cell aging: Regulation and rejuvenation. Trends in Endocrinology and Metabolism 26, 287–296.

Subauste, M.C., Pertz, O., Adamson, E.D., Turner, C.E., Junger, S., and Hahn, K.M. (2004). Vinculin modulation of paxillin-FAK interactions regulates ERK to control survival and motility. Journal of Cell Biology 165, 371–381.

Susaki, E.A., Tainaka, K., Perrin, D., Kishino, F., Tawara, T., Watanabe, T.M., Yokoyama, C., Onoe, H., Eguchi, M., Yamaguchi, S., et al. (2014). Whole-brain imaging with single-cell resolution using chemical cocktails and computational analysis. Cell 157, 726–739.

Suzuki, J., Yamazaki, Y., Li, G., Kaziro, Y., and Koide, H. (2000). Involvement of Ras and Ral in Chemotactic Migration of Skeletal Myoblasts. Molecular and Cellular Biology 20, 7049–7049.

Takaoka, S., Kamioka, Y., Takakura, K., Baba, A., Shime, H., Seya, T., and Matsuda, M. (2016). Live imaging of transforming growth factor-β activated kinase 1 activation in Lewis lung carcinoma 3LL cells implanted into syngeneic mice and treated with polyinosinic: Polycytidylic acid. Cancer Science 107, 644–652.

Tedesco, F.S., Dellavalle, A., Diaz-Manera, J., Messina, G., and Cossu, G. (2010). Repairing skeletal muscle: regenerative potential of skeletal muscle stem cells. J Clin Invest 120, 11–19.

Teranishi, S., Kimura, K., and Nishida, T. (2009). Role of formation of an ERK-FAK-paxillin complex in migration of human corneal epithelial cells during wound closure in vitro. Investigative Ophthalmology and Visual Science 50, 5646–5652.

Tinevez, J.Y., Perry, N., Schindelin, J., Hoopes, G.M., Reynolds, G.D., Laplantine, E., Bednarek, S.Y., Shorte, S.L., and Eliceiri, K.W. (2017). TrackMate: An open and extensible platform for single-particle tracking. Methods 115, 80–90.

Tintignac, L.A.J., Sirri, V., Leibovitch, M.P., Lecluse, Y., Castedo, M., Metivier, D., Kroemer, G., and Leibovitch, S.A. (2004). Mutant MyoD Lacking Cdc2 Phosphorylation Sites Delays M-Phase Entry. Molecular and Cellular Biology 24, 1809–1821.

Tomura, M., Sakaue-Sawano, A., Mori, Y., Takase-Utsugi, M., Hata, A., Ohtawa, K., Kanagawa, O., and Miyawaki, A. (2013). Contrasting Quiescent G0 Phase with Mitotic Cell Cycling in the Mouse Immune System. PLoS ONE 8, 1–10.

Webster, M.T., Manor, U., Lippincott-Schwartz, J., and Fan, C.M. (2016). Intravital Imaging Reveals Ghost Fibers as Architectural Units Guiding Myogenic Progenitors during Regeneration. Cell Stem Cell 18, 243–252.

Yano, S., Miwa, S., Mii, S., Hiroshima, Y., Uehara, F., Yamamoto, M., Kishimoto, H., Tazawa, H., Bouvet, M., Fujiwara, T., et al. (2014). Invading cancer cells are predominantly in G0/G1 resulting in chemoresistance demonstrated by real-time FUCCI imaging. Cell Cycle 13, 953–960.

Yin, H., Price, F., and Rudnicki, M.A. (2013). Satellite Cells and the Muscle Stem Cell Niche. Physiological Reviews 93, 23–67.

Yokoyama, T., Takano, K., Yoshida, A., Katada, F., Sun, P., Takenawa, T., Andoh, T., and Endo, T. (2007). DA-Raf1, a competent intrinsic dominant-negative antagonist of the Ras-ERK pathway, is required for myogenic differentiation. Journal of Cell Biology 177, 781–793.

